# An fMRI dataset of verbalized spontaneous thought with annotated transcripts and self-report trait measures

**DOI:** 10.64898/2026.05.12.724488

**Authors:** Mengting Zhang, Philip R. Liu, Haowen Su, Mengqi Zhao, Xian Li, Savannah Born, Yoonjung Lee, Christopher J. Honey, Janice Chen, Hongmi Lee

## Abstract

Spontaneous thought is pervasive in everyday human cognition, yet datasets capturing its neural dynamics under minimally interrupted conditions remain limited. The current dataset was acquired from a think-aloud functional MRI experiment in which 118 participants continuously verbalized their spontaneous thoughts during 10-minute scanning sessions. The raw MRI data and verbal transcripts with sentence-level timestamps were previously released and analyzed in our prior study examining neural activity associated with thought transitions. Building on that release, we additionally provide preprocessed MRI data, speech transcriptions with word-level timestamps aligned to image acquisition, large language model-generated ratings of transcribed thoughts across emotional and sensory dimensions, and self-report survey measures assessing personality, mental health, and cognitive abilities. Validation analyses demonstrated activation in expected cortical regions associated with speech production and sensory content identified from transcript annotations, agreement between language model and human ratings, and adequate internal consistency of survey measures, supporting the dataset’s overall quality. This dataset enables reuse for investigations of spontaneous thought, speech generation, and individual differences using naturalistic functional MRI data.

## BACKGROUND & SUMMARY

Humans frequently experience spontaneous thoughts that arise without explicit goals or direct responses to external stimuli^1–5^. These internally generated thoughts are fundamental to everyday cognition, supporting autobiographical memory, future planning, creativity, and self-reflection^6–8^. Alterations in spontaneous thought patterns, such as persistent rumination or intrusive thoughts, have also been implicated in psychiatric conditions including depression, anxiety, and post-traumatic stress disorder (PTSD)^2,9,10^. Accordingly, extensive research has employed diverse experimental paradigms to investigate the cognitive and neural mechanisms underlying spontaneous thought^11–16^.

Among these approaches, the think-aloud paradigm has recently received renewed attention^17–20^. Originally developed to study problem solving and decision making^21,22^, this method instructs participants to continuously verbalize their thoughts as they occur. Empirical studies have shown that verbalizing thoughts aloud does not meaningfully alter their phenomenological qualities or content relative to silent thinking, supporting the validity of this paradigm for studying spontaneous thought during rest^23,24^. Moreover, compared with traditional methods such as intermittent experience sampling^25–27^ or retrospective self-report^28,29^, the think-aloud paradigm provides more direct access to the real-time, continuous temporal dynamics of thought content while preserving the natural flow of consciousness^18,30,31^. Leveraging this advantage, prior research has examined the structure of ongoing thought streams, demonstrating that thoughts typically form clusters of semantically related content before transitioning to new topics^17,19^. The variability and stability of these thought trajectories have been associated with different states of cognitive control^32^ and are thought to contribute to emotion regulation^20^.

Despite its widespread use in behavioral research, the think-aloud paradigm has rarely been combined with neuroimaging to examine the neural dynamics underlying spontaneous thought. One of the few neuroimaging studies^30^ used functional magnetic resonance imaging (fMRI) during a think-aloud task and, through natural language processing and representational similarity analysis^33^, showed that representations of thought content are distributed across multiple large-scale brain networks. A more recent study^34^ applied latent state modeling to think-aloud fMRI data to identify recurring brain states associated with thought orientation (internal versus external) and novelty. Although these studies released datasets containing raw or processed MRI data along with selected participant- or thought-level features^35,36^, neither study made the raw think-aloud transcripts publicly available, thereby limiting opportunities for data reuse and content-level analyses.

Here, we release an extended version of our previously published think-aloud fMRI dataset, which originally included raw MRI data and sentence-level annotated transcripts^31,37,38^. The dataset was collected from 118 participants, each of whom completed a 10-minute think-aloud session while undergoing MRI scanning. Participants’ verbal responses were transcribed, segmented and timestamped at the sentence level, and manually annotated for thought category (e.g., episodic memory, future thinking) and topic. Analyses of the original dataset showed that transitions between thoughts, especially those involving changes in topic rather than thought category, engage regions within the brain’s default mode^39,40^ and control networks^41^, producing activation patterns resembling those observed at boundaries between external events^42^. In addition, functional connectivity within and between these networks predicted the semantic variability of individuals’ thought trajectories.

In the current data descriptor, we substantially extend the previously shared dataset by additionally providing: (1) standardized fMRIPrep^43^-preprocessed MRI data with quality control metrics, (2) fine-grained word-level speech timestamps aligned to fMRI acquisition, (3) sensory and affective ratings of sentence-level thought content generated using a large language model (LLM) and validated against human annotations, and (4) an extensive set of post-scan survey measures assessing personality, mental health, and cognitive abilities.

These additional features support a broad range of uses. First, the inclusion of standardized preprocessed outputs minimizes preprocessing time and effort, thereby lowering technical barriers and facilitating reuse by researchers with diverse levels of neuroimaging expertise. Second, word-level speech timestamps offer substantially finer temporal resolution than sentence-level segmentation, enabling more precise analyses of neural activity associated with specific linguistic features and semantic content during speech production. Third, the LLM-generated multidimensional sensory and affective ratings of transcribed thoughts support analyses of neural responses associated with the perceptual and emotional aspects of spontaneous thought. LLMs have been shown to perform comparably to human annotators across various text annotation tasks^44,45^ and have been successfully applied in prior work to quantify multiple psychological dimensions of thought content^34,46,47^. Finally, the broad set of survey measures allows investigation of relationships between spontaneous thought patterns and a wide range of trait-level variables in individual differences analyses. Previous studies have linked think-aloud language to personality traits^46,48^ as well as mental health characteristics^18,49,50^. The current dataset provides an opportunity to relate these behavioral findings to their underlying neural mechanisms.

## METHODS

The study procedures complied with ethical standards for research involving human participants as outlined in the Declaration of Helsinki. All procedures were conducted in accordance with protocols approved by the Institutional Review Board (IRB) of Johns Hopkins Medicine (approval number: IRB00201118).

### Participants

A total of 126 healthy adults (aged 18–40 years; M = 23.7) were recruited from the Johns Hopkins University community. All participants were right-handed native English speakers with normal hearing and normal or corrected-to-normal vision. Informed consent was obtained in accordance with procedures approved by the Johns Hopkins Medicine IRB.

Eight participants were excluded from both the MRI and transcript datasets for the following reasons: poor speech audio recording quality (N = 5), scanning interruptions due to technical issues (N = 2), and failure to adhere to task instructions (N = 1). After these exclusions, data from 118 participants (73 females; aged 18–39 years; M = 23.4 years) were included in the MRI and transcript datasets and shared publicly.

### Experimental task and procedures

Participants completed a 10-minute think-aloud session during fMRI scanning, during which they were instructed to continuously verbalize their spontaneous stream of thoughts (Fig. 1a). They were encouraged to speak freely about whatever came to mind, such as memories, plans, and current perceptual experiences, while avoiding elaboration or explanation for the experimenter.

**Fig. 1.**
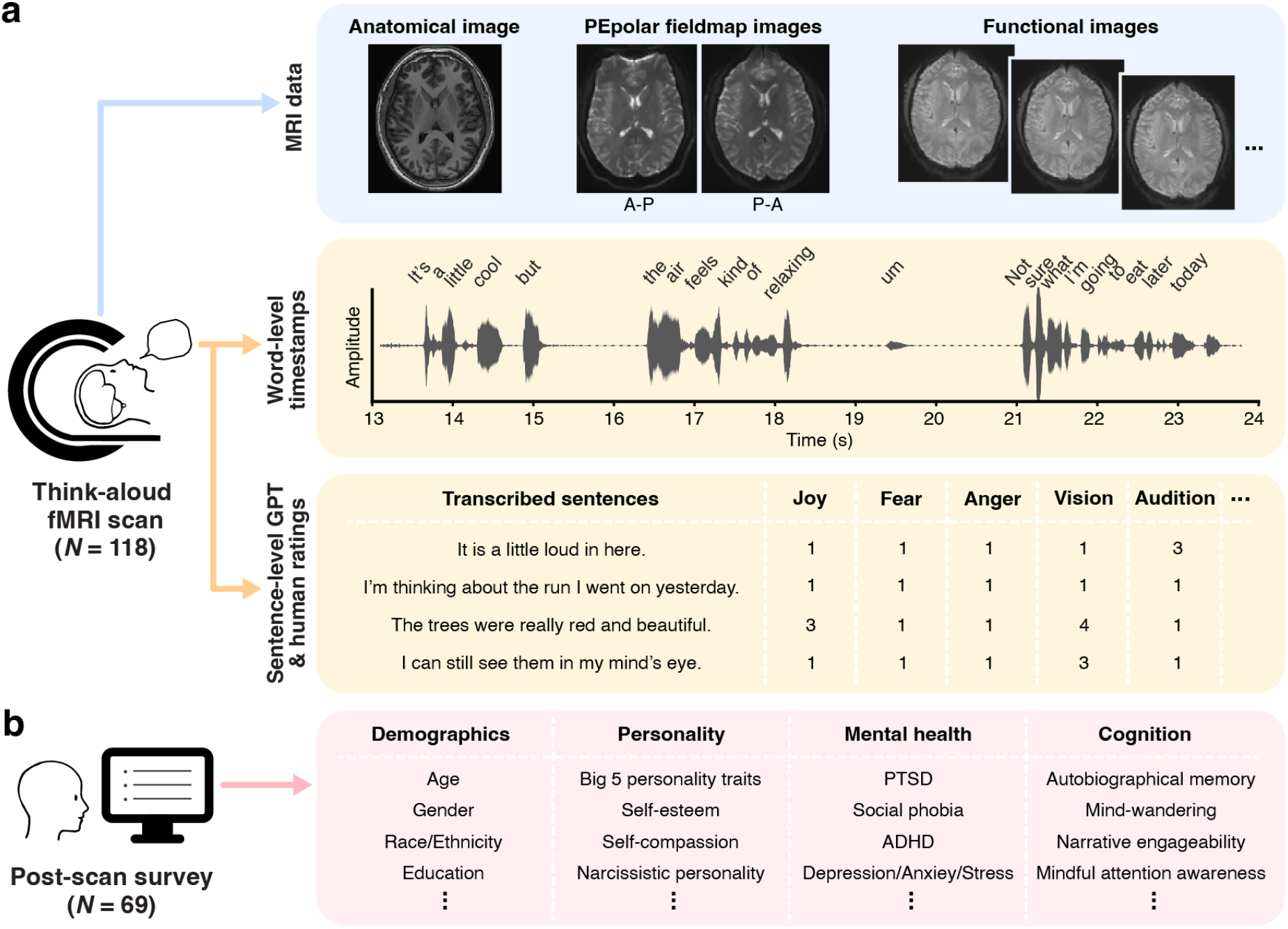
Overview of the dataset. (**a**) During fMRI scanning, participants completed a continuous think-aloud task in which they verbally reported their spontaneous stream of thoughts. The upper panel depicts the neuroimaging data included in the dataset, comprising anatomical (T1-weighted) images, fieldmap images acquired with anterior–posterior (A–P) and posterior–anterior (P–A) phase-encoding directions, and functional images. Representative slices from a single participant are shown for illustration. The middle panel illustrates word-level speech timestamps derived from audio recordings of participants’ verbal responses. The lower panel shows sentence-level segments of these responses, which were evaluated for affective properties (e.g., emotional intensity and discrete emotions) and sensory content (e.g., visual, auditory) by both a large language model (specifically, Generative Pre-trained Transformer 5; GPT-5) and human raters. Selected rating dimensions and example scores are displayed for illustration. (**b**) A subset of the fMRI participants completed online surveys within two days of the scan, assessing demographic characteristics, personality traits, mental health, and cognitive abilities. PTSD = post-traumatic stress disorder, ADHD = attention-deficit/hyperactivity disorder.

The task began with a brief “Begin” cue, followed by a fixation cross presented on a gray background for the remainder of the session. The visual stimuli were projected onto a screen at the rear of the scanner bore using Psychophysics Toolbox Version 3 (http://psychtoolbox.org). Participants’ speech was recorded with an MR-compatible microphone (FOMRI II; Optoacoustics Ltd.). Participants were asked to minimize head movement by speaking primarily with their jaw while keeping their head still. A detailed description of the experimental setup and task instructions is provided in ref.^31^.

### MRI data acquisition

MRI data were collected at the Kennedy Krieger Institute (Baltimore, MD) on a 3 T Philips Ingenia Elition scanner using a 32-channel head coil. Functional images were acquired with a T2*-weighted multiband echo-planar imaging (EPI) sequence (TR = 1.5 s, TE = 30 ms, flip angle = 52°, acceleration factor = 4, voxel size = 2 × 2 × 2 mm^3^). Fieldmap (2 × 2 × 2 mm^3^) and high-resolution T1-weighted anatomical images (1 × 1 × 1 mm^3^) were also collected. Additional acquisition details can be found in ref.^31^.

### MRI data preprocessing

The original MRI data from our published study^31^ were made publicly available via OpenNeuro^38^. All MRI data were provided in NIfTI format and organized in accordance with the Brain Imaging Data Structure (BIDS) specification^51^. For public release, anatomical images were defaced using pydeface (version 2.0.2) to remove identifiable facial features and protect participant privacy.

Preprocessing of the raw (non-defaced) anatomical images and functional images was performed using fMRIPrep^43^ (version 23.2.1) with default settings, except that slice timing correction was not applied. Anatomical images were corrected for intensity nonuniformity, skull-stripped, and segmented into cerebrospinal fluid (CSF), white matter, and gray matter. Volume-based spatial normalization to a standard template (MNI152NLin2009cAsym) was performed using nonlinear registration. In addition, cortical surface reconstruction was carried out using FreeSurfer^52^ (version 7.3.2). Functional images were corrected for head motion and magnetic field inhomogeneities and subsequently normalized to both the MNI152NLin2009cAsym template in volume space and the FreeSurfer fsaverage6 surface space. Magnetic field inhomogeneity correction was not applied for one participant due to an error in fieldmap processing. This is documented in the MRI dataset metadata (readme.txt).

Following fMRIPrep preprocessing, functional images in volume space were spatially smoothed with a 4 mm full-width-at-half-maximum (FWHM) Gaussian kernel using the NLTools Python package (version 0.5.1). Residual head motion and physiological noise were further reduced by regressing out the six motion parameters (three translations and three rotations) estimated during fMRIPrep motion correction, CSF and white matter signals, and second-order polynomial trends. The resulting residual time series were subsequently z-scored along the temporal dimension. All metadata and output files associated with MRI preprocessing, including the regressors used for the noise-reduction step, are available in the MRI dataset (see Data Records below).

### Transcription and timestamp alignment

Audio recordings of participants’ think-aloud responses were either manually transcribed or automatically transcribed using Whisper (Large-v2 model; OpenAI) and subsequently verified and corrected by human annotators. Each transcript was segmented into individual sentences, with timestamps identified for the onset and offset of each sentence, as described in detail in ref.^31^. The original sentence-segmented transcript data, along with thought category and topic annotations, were made publicly available via Zenodo^37^.

To obtain finer-grained temporal alignment, we further generated word-level timestamps using Gentle (version 0.11.0; https://github.com/strob/gentle), an open-source forced-alignment tool that maps each word in the verified transcript to its corresponding segment in the audio recording. Prior to alignment, transcripts were concatenated across sentences and preprocessed to improve matching accuracy by converting numerals and time expressions into words and removing special punctuation. Each participant’s audio file, together with the preprocessed transcript in text-file format, was provided as input to Gentle, which generated onset and offset timestamps for individual words in seconds relative to the start of the audio recording. These timestamps were adjusted so that time zero corresponded to the onset of the first volume acquisition. The resulting output was converted into a spreadsheet containing three columns—word, start time, and end time—with missing values for words that could not be automatically localized. These files were subsequently manually reviewed and revised by twelve trained research assistants. They listened to the audio while inspecting the existing timestamps to add timestamps for words that were not successfully aligned and to correct machine-generated errors, including overlapping or inverted time intervals. They also corrected transcription errors and added filler words (e.g., uh, um) that were missing from the original transcripts.

For 54 of the 118 transcripts with relatively poor audio quality, we additionally generated word-level timestamps using WhisperX (version 3.7.4; OpenAI), which also provides automatic word-level alignment from the audio signal. WhisperX timestamps were used as a reference to fill missing timestamps in the Gentle output. The trained research assistants manually integrated the Gentle and WhisperX outputs and reviewed the combined files following the same procedure described above.

### Psychological dimension ratings

We extended the previously published annotation of the think-aloud transcripts^37^ by providing ratings for each transcript segment (sentence) across 14 psychological dimensions capturing both affective and sensory aspects of the reported thoughts. Affective dimensions included emotional intensity, joy, sadness, fear, anger, disgust, surprise, and anxiety. Emotional intensity was rated from −4 (very negative) to +4 (very positive), with neutral states rated close to zero. The remaining affective dimensions were rated from 1 (not at all) to 4 (very much), reflecting the degree to which each emotion appeared to be experienced in the described thought. Sensory dimensions assessed the extent to which perceptual or bodily experiences were present in each thought, including vision, audition, olfaction, gustation, somatosensation, and interoception, each rated on a 1−4 scale.

A large language model (LLM), Generative Pre-trained Transformer 5 (GPT-5; Nano model; OpenAI), was used to automatically generate these ratings using few-shot prompting with default settings (medium verbosity and reasoning effort; no sampling parameters specified). The model was guided by a standardized prompt describing the task context, which was applied consistently across all transcripts. Specifically, the model was informed that it should rate individual thoughts spoken aloud by a participant during an fMRI experiment and evaluate each thought along a specified psychological dimension based solely on the content of the current thought using the provided rating scales. Ratings were generated using a dimension-by-dimension approach, in which the model rated all thoughts within a transcript for one dimension before proceeding to the next. For each dimension, example sentences paired with corresponding ratings were provided as few-shot demonstrations (e.g., emotional intensity for “winning that award felt like the best moment of my life so far”: 4). Each rating was generated once without repetition. The exact prompts, including the task description and dimension-specific examples, are available in the shared code repository within the OSF dataset (code/llm_rating/config/examples_lite.json; code/llm_rating/config/prompts/system_prompt.txt and dimension_wise_prompt.txt).

To validate the LLM-generated ratings, we additionally collected corresponding human ratings for a randomly selected subset of 18 transcripts, evaluated by four independent human raters. To reduce rater fatigue, transcripts were sampled from those with fewer sentences than the sample median (97 sentences). Raters were trained using the same instructions and example materials provided to the LLM. Consistent with the LLM procedure, human raters also completed evaluations in a dimension-by-dimension manner across thoughts and were instructed to apply consistent rating criteria within each dimension and across transcripts.

### Post-scan survey

Of the 126 participants recruited for the fMRI session, 69 completed and submitted an online battery of self-report surveys administered via Qualtrics (Fig. 1b). Participants were instructed to complete the surveys within two days of their scanning session. The survey battery assessed a broad range of individual differences spanning personality traits, mental health, and cognitive abilities, as detailed below.

The Big Five Inventory^53^ is a 44-item questionnaire that measures five major dimensions of personality: Openness, Conscientiousness, Extraversion, Agreeableness, and Neuroticism. Each item is rated on a 5-point Likert scale ranging from 1 (strongly disagree) to 5 (strongly agree). Higher total scores within each personality dimension indicate a stronger tendency toward that dimension.

The Survey of Autobiographical Memory (SAM)^54^ measures individual differences in autobiographical memory across four domains: episodic, semantic, spatial, and future-oriented memory. The episodic domain reflects the vividness and richness of personal memories, the semantic domain captures access to factual and self-related knowledge, the spatial domain assesses one’s ability to recall and navigate environments, and the future domain measures the capacity to imagine or simulate future events. Participants rate each statement on a 5-point Likert scale ranging from 1 (strongly disagree) to 5 (strongly agree), with higher total scores indicating stronger self-reported ability within that domain.

The Mind-Wandering Questionnaire (MWQ)^55^ is a 5-item measure that assesses the frequency of unintentional mind-wandering in daily life. Each item is rated on a 6-point Likert scale ranging from 1 (almost never) to 6 (almost always), with higher total scores indicating a greater tendency for thoughts to drift away from the present moment.

The Mindful Attention Awareness Scale (MAAS)^56^ is a 15-item measure of dispositional mindfulness, reflecting awareness and attention to present-moment experiences. Each item is rated on a 6-point Likert scale ranging from 1 (almost always) to 6 (almost never), with higher average scores indicating greater mindfulness and attentional awareness.

The White Bear Suppression Inventory (WBSI)^57^ is a 15-item measure that assesses the general tendency to suppress unwanted thoughts. Each item is rated on a 5-point Likert scale ranging from 1 (strongly disagree) to 5 (strongly agree), with higher total scores indicating a stronger tendency toward thought suppression.

The Automatic Thoughts Questionnaire (ATQ-30)^58^ captured the frequency of negative self-referential cognitions associated with depressive thinking. Participants indicated how often they experienced each of 30 statements on a 5-point scale (1 = not at all, 5 = all the time); higher total scores denote more frequent negative automatic thoughts.

The Adult ADHD Self-Report Scale (ASRS-v1.1)^59^ is a brief 6-item measure that assesses symptoms of attention-deficit/hyperactivity disorder in adults. Each item is rated on a 5-point scale ranging from 1 (never) to 5 (very often), with higher total scores indicating greater symptom frequency.

The Narcissistic Personality Inventory-40 (NPI-40)^60^ assesses non-clinical expressions of narcissism through 40 paired statements. For each pair, participants choose the statement that best describes them, capturing traits such as authority, entitlement, exhibitionism, exploitativeness, self-sufficiency, superiority, and vanity. Selecting the narcissistic statement is scored as 1, whereas selecting the non-narcissistic statement is scored as 0. Higher total scores indicate greater endorsement of narcissistic traits.

The Rosenberg Self-Esteem Scale (RSES)^61^ measures global self-worth through 10 statements rated on a 4-point scale ranging from 1 (strongly disagree) to 4 (strongly agree). Higher total scores indicate greater self-esteem and a more positive self-evaluation.

The Self-Compassion Scale–Short Form (SCS-SF)^62^ includes 12 items that assess an individual’s tendency to respond to personal challenges with self-kindness, mindfulness, and a sense of shared humanity. Each item is rated on a 5-point scale ranging from 1 (almost never) to 5 (almost always), and higher averages indicate greater self-compassion.

The Brief Selfism Scale^63^ consists of 6 items that assess general attitudes of self-interest and self-prioritization, selected from the 28-item Selfism Scale^64^. Participants rated the extent to which each statement described them on a 5-point scale ranging from 1 (strongly disagree) to 5 (strongly agree). Higher total scores indicate stronger endorsement of self-focused or self-serving attitudes.

The Curiosity and Exploration Inventory-II (CEI-II)^65^ assesses trait curiosity across two dimensions: Stretching, which reflects the motivation to seek new knowledge and experiences, and Embracing, which reflects openness to uncertainty and novelty. The scale includes 10 items rated on a 5-point scale ranging from 1 (very slightly or not at all) to 5 (extremely), with higher total scores indicating greater curiosity and exploratory drive.

The Depression Anxiety Stress Scales-21 (DASS-21)^66^ is a 21-item measure that assesses emotional distress across three dimensions: depression, anxiety, and stress. Each item is rated on a 4-point scale ranging from 0 (did not apply to me at all) to 3 (applied to me very much or most of the time). Scores were summarized separately for each dimension, with higher total scores indicating greater symptom severity.

The PTSD Checklist for DSM-5 (PCL-5)^67^ measures post-traumatic stress symptoms across 20 items rated on a 5-point scale ranging from 1 (not at all) to 5 (extremely). Total scores provide an index of overall PTSD symptom severity, with higher scores indicating more frequent or intense symptoms.

The State-Trait Inventory for Cognitive and Somatic Anxiety – Trait version (STICSA)^68^ assesses stable individual differences in anxiety through 21 items rated on a 4-point scale ranging from 1 (not at all) to 4 (very much so). Higher total scores indicate greater levels of trait anxiety.

The Social Phobia Inventory (SPIN)^69^ measures symptoms of social anxiety, including fear, avoidance, and physiological discomfort in social situations. The scale consists of 17 items rated on a 5-point scale ranging from 1 (not at all) to 5 (extremely), and higher total scores indicate more severe social anxiety.

The Ruminative Responses Scale-10 (RRS-10)^70^ assesses the tendency to engage in repetitive, self-focused thinking following negative mood states. The scale includes 10 items that reflect two conceptual components, Brooding and Reflection. Each item is rated on a 4-point scale ranging from 1 (almost never) to 4 (almost always), with higher total scores indicating a stronger tendency to ruminate.

The Narrative Engageability Scale^71^ measures a trait-level tendency to become cognitively and emotionally immersed in stories. It consists of 16 items rated on a 7-point scale ranging from 1 (strongly disagree) to 7 (strongly agree). Higher average scores indicate greater narrative engagement and empathy during story processing.

After completing all surveys, participants provided demographic information. They reported their age, gender, race/ethnicity, Hispanic identity, sexual orientation, highest level of education, annual household income, country of birth, country of citizenship, religious affiliation, and political orientation. In addition, participants rated how comfortable they felt describing their thoughts during the think-aloud fMRI session on a 5-point scale (ranging from extremely uncomfortable to extremely comfortable) and indicated the extent to which they censored themselves while speaking on a 3-point scale (not at all, somewhat, very much).

### DATA RECORDS

The dataset is publicly available across two repositories. Neuroimaging data can be accessed through the OpenNeuro repository (accession number: ds006067; version 2.0.0)^72^. Behavioral data including think-aloud transcripts with timestamps, sentence-level psychological dimension ratings, and post-scan survey responses are available via the Open Science Framework (OSF) under the project titled “Think Aloud Behavioral Data”^73^. Code for data generation, preprocessing, and validation analyses is also available in the same OSF repository.

### MRI data

Raw neuroimaging data for this study were previously released on OpenNeuro (version 1.0.1)^38^ and are described in ref.^31^. The dataset includes anatomical, field map, and functional MRI scans for 118 participants, organized in a BIDS-compliant format^51^.

To facilitate data reuse, we extended the previously released dataset by additionally providing images preprocessed using the fMRIPrep and FreeSurfer pipelines, along with their associated output files and all necessary metadata. fMRIPrep preprocessing outputs are organized within the “derivatives” directory and grouped by participant (sub-<ID>), following BIDS-Derivatives conventions. FreeSurfer outputs are located in the “derivatives/sourcedata/freesurfer” directory and organized by participant (sub-<ID>), following the standard FreeSurfer recon-all pipeline structure.

A detailed description of preprocessing outputs is available in the fMRIPrep documentation (https://fmriprep.org/en/stable/outputs.html). The fMRIPrep-preprocessed anatomical image in volume space (sub-<ID>_space-MNI152NLin2009cAsym_desc-preproc_T1w.nii.gz) can be found in the “anat” directory of each participant’s folder. The fMRIPrep-preprocessed functional image in volume space (sub-<ID>_task-thinkaloud_space-MNI152NLin2009cAsym_desc-preproc_bold.nii.gz), as well as the additionally smoothed, noise-reduced, and z-scored image (sub-<ID>_task-thinkaloud_space-MNI152NLin2009cAsym_desc-preproc_smooth4mm_denoise_bold.nii.gz), are available in the “func” directory of each participant’s folder. Confound regressors, including the variables used in the additional noise-reduction step, are also provided as tab-separated value files (sub-<ID>_task-thinkaloud_desc-confounds_timeseries.tsv) in the same “func” directory.

### Transcripts with timestamps

Think-aloud transcript files with word-level timestamps are available in the “data/transcripts_and_timestamps/word_level” directory in the OSF repository. The files are provided in Excel spreadsheet format (sub-<ID>_timestamps.xlsx), with one row per spoken word and three columns: Transcribed Word, Start Time, and End Time.

For completeness, we also provide the sentence-level transcripts previously released alongside our published study^31^, alongside the newly released data. The sentence-level transcript files are also available in Excel spreadsheet format (sub-<ID>_transcripts.xlsx) in the “data/transcripts_and_timestamps/sentence_level” directory. Each file contains one row per spoken sentence and includes the following columns: Transcribed Sentence, Start Time, and End Time. For both word- and sentence-level data, timestamps are specified in seconds relative to the acquisition of the first volume of the functional image.

### Sentence-level ratings

GPT-generated ratings for the 14 psychological dimensions corresponding to each transcribed sentence are available in the “data/sentence_level_ratings/gpt_generated” directory of the OSF repository. The rating files are provided in Excel spreadsheet format (sub-<ID>.xlsx), with each row corresponding to a single sentence. Each file contains 15 columns, including the Transcribed Sentence column and numerical ratings for all 14 psychological dimensions.

Ratings for the 18 transcripts independently annotated by four human raters are available in the “data/sentence_level_ratings/human_validation” directory, organized by rater. Each rater-specific folder contains 18 Excel files (sub-<ID>.xlsx), with each file including the same columns as the GPT-generated rating files. Rater identities are anonymized, and no personally identifiable information is included.

### Post-scan survey

Post-scan survey data files are available in the “data/questionnaires” directory of the OSF repository. The original survey materials are provided as a Word document (Think_Aloud_Personality_Surveys.docx), which contains all items from the full set of questionnaires administered in the study. Item-level responses are provided in ThinkAloudQuestionnaires_preproc.csv. This file contains participants’ raw responses to demographic questions and individual questionnaire items, with columns labeled by survey name and item number. Individual-level summary scores are provided in ThinkAloudQuestionnaires_summary.csv, which contains demographic variables and composite questionnaire scores computed from item-level responses according to each instrument’s scoring procedures. In both the item-level and summary data files, each row corresponds to a single participant, and the BIDS_ID column indicates the subject identifier used to link the survey data with the transcript and MRI datasets.

## TECHNICAL VALIDATION

We conducted a series of validation analyses to assess the quality and validity of each component of the dataset, including anatomical and functional neuroimaging data, word-level speech timestamps, LLM-generated content ratings, and self-report survey measures.

### Anatomical neuroimaging data quality

We extracted quality metrics for anatomical MRI data (Fig. 2a) using the MRI Quality Control tool (MRIQC; version 22.0.6)^74^, a widely used software package for automated MRI quality assessment. MRIQC applies minimal preprocessing (e.g., skull-stripping, tissue segmentation) to compute standardized quality control metrics comparable across studies.

**Fig. 2.**
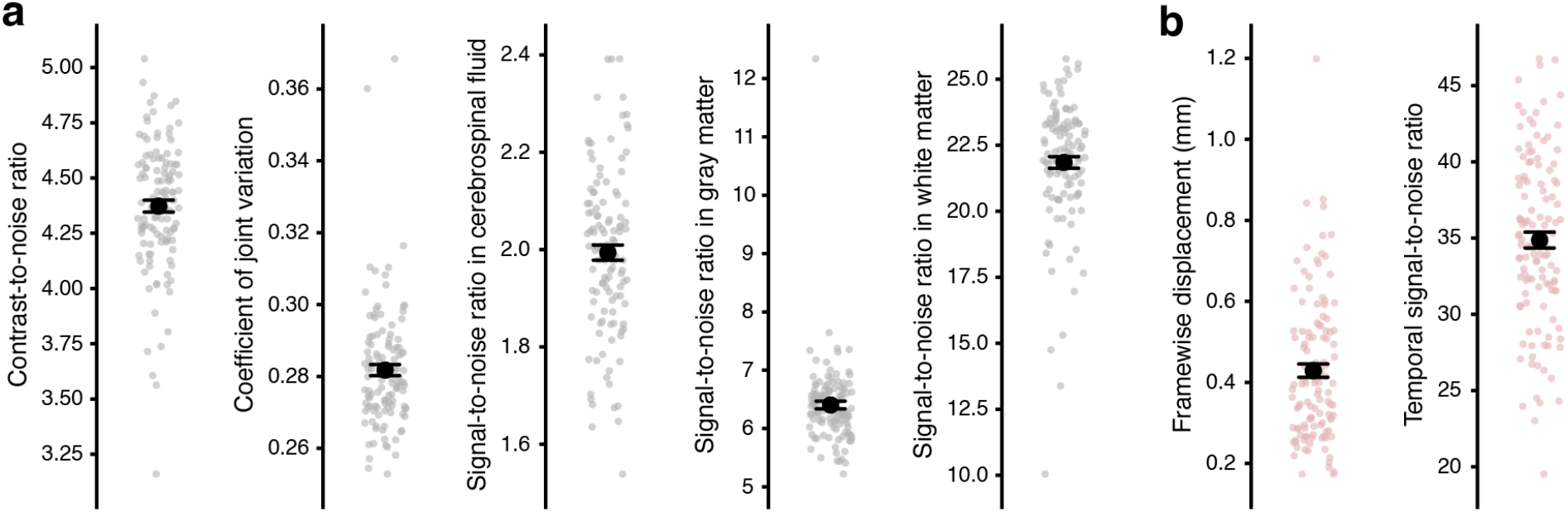
MRI data quality metrics. (**a**) Anatomical MRI quality metrics, including contrast-to-noise ratio, coefficient of joint variation, and signal-to-noise ratio in cerebrospinal fluid, gray matter, and white matter. (**b**) Functional MRI quality metrics, including mean framewise displacement and temporal signal-to-noise ratio. In both panels, individual participant values are shown as small dots (N = 118). Larger black circles indicate the mean across participants, and error bars represent 95% bootstrap confidence intervals.

The contrast-to-noise ratio (CNR) was calculated as the difference in mean intensity between gray matter and white matter divided by the noise level, reflecting the ability to distinguish these tissue types^75^. The average CNR across participants was 4.37 (SD = 0.29). We additionally computed the coefficient of joint variation (CJV), which quantifies variability between gray matter and white matter intensities and is sensitive to intensity inhomogeneity and motion artifacts; lower CJV values indicate better image quality^76^. The mean CJV was 0.28 (SD = 0.017). Signal-to-noise ratio (SNR) was defined as the mean intensity within each tissue type divided by the standard deviation of background noise^75^. The mean SNR values for CSF, gray matter, and white matter were 1.99 (SD = 0.17), 6.40 (SD = 0.72), and 21.84 (SD = 2.4), respectively. These quality metric values are comparable to those reported in previously published datasets^77,78^, suggesting adequate structural image quality across participants.

In addition, all anatomical images were visually inspected by the authors. Two participants showed anatomical anomalies (enlarged ventricles). These cases are documented in the MRI dataset metadata (readme.txt) and are included in all quality metric summaries to provide a comprehensive overview of data quality across the full sample.

### Functional neuroimaging data quality

For functional MRI data quality assessment (Fig. 2b), we first examined framewise displacement (FD), which summarizes volume-to-volume changes in head position. FD metrics were derived from the fMRIPrep^43^ preprocessing outputs. Across the 10-minute think-aloud session, mean FD averaged 0.43 mm (SD = 0.18; range = 0.17-1.20 mm) across participants. This level of motion is generally higher than that observed in typical video-watching scans (e.g., refs.^77,79,80^), reflecting the demands of the task which required participants to speak aloud during scanning.

To evaluate overall data quality following preprocessing, we additionally calculated the temporal signal-to-noise ratio (tSNR), a commonly used metric for assessing sensitivity to detect brain activation in fMRI data^81^. tSNR was computed from the functional images output by fMRIPrep in volume space, prior to the additional preprocessing steps, including spatial smoothing, temporal filtering, and z-scoring. For each voxel, tSNR was defined as the mean blood-oxygenation-level-dependent (BOLD) signal across time divided by its temporal standard deviation. Voxelwise tSNR values were then summarized at the participant level as the median tSNR across all gray matter voxels.

Median tSNR values averaged 34.87 across participants (SD = 5.65; range = 19.52-46.76). Because tSNR is influenced by various imaging parameters, strict absolute thresholds are difficult to define, and values should be interpreted in the context of the specific acquisition protocol^82^. That said, tSNR values in the current dataset were generally lower than those reported in other naturalistic fMRI studies^79,80,83^, again likely reflecting increased head motion, a well-established source of signal instability in fMRI^84^. To directly assess this possibility, we examined the relationship between mean FD and median tSNR across participants and observed a significant negative Pearson correlation between the two (*r*(116) = -0.79, *p* < 0.001). For fMRI analyses in our published study^31^ and in the validation analyses reported below, participants with excessive head motion (mean FD > 0.5 mm) were excluded. We similarly recommend that dataset users consider applying motion-based exclusion criteria in their analyses.

### Word-level timestamp validation

We first examined descriptive statistics of the word-level timestamps derived from the think-aloud transcripts. Participants produced an average of 1395.5 words (SD = 382, range: 381– 2280) during the 10-minute session, corresponding to approximately 139.5 words per minute, consistent with typical English conversational speech rates^85,86^. The mean word duration was 233.5 ms (SD = 31.12).

To further validate the utility of the word-level timestamps for neuroimaging analyses, we examined brain activation associated with overt speech production as identified from these timestamps. If the timestamps reliably capture periods of speech, regions involved in speech motor control and auditory processing should show greater activation during speech than during silence. Within each participant, speech periods were defined as TRs containing at least one word onset, whereas silence periods were defined as TRs with no detected speech. To minimize contamination from carryover speech-related activity during brief pauses, only silence periods consisting of at least 3 consecutive TRs (4.5 s) were included in the analysis. To account for hemodynamic response delay, all timestamps were shifted forward by 4.5 s relative to the corresponding speech events.

A total of 73 participants were included in the analysis after excluding 39 participants due to excessive motion, 2 due to anatomical anomalies, 1 due to technical issues during scanning, 1 due to MRI artifacts, and 2 due to the absence of sufficiently long silence periods. For each participant and cortical parcel defined by the Schaefer 400-parcel whole-brain cortical atlas^87^, preprocessed BOLD signals were averaged across TRs separately for speech and silence periods, and a speech-versus-silence contrast was computed. Motion outlier TRs (FD ≥ 1 mm), along with the two immediately preceding and following TRs, were excluded from analysis. Group-level effects were then assessed across participants using two-tailed one-sample *t*-tests against zero on these within-subject contrasts. Multiple comparisons correction across parcels was applied using the false discovery rate (FDR) procedure^88^ (*q* < 0.05).

The resulting whole-brain group-level contrast maps (Fig. 3) revealed activation during speech periods relative to silence periods across a broad network of cortical regions. Significant effects were observed in the bilateral somatomotor cortex, frontal opercular regions, superior temporal cortex, and medial frontal areas. These regions have been consistently implicated in motor execution, articulation, and auditory monitoring during overt speech production^30,89,90^, supporting the validity of the word-level timestamps for capturing speech-related neural activity.

**Fig. 3.**
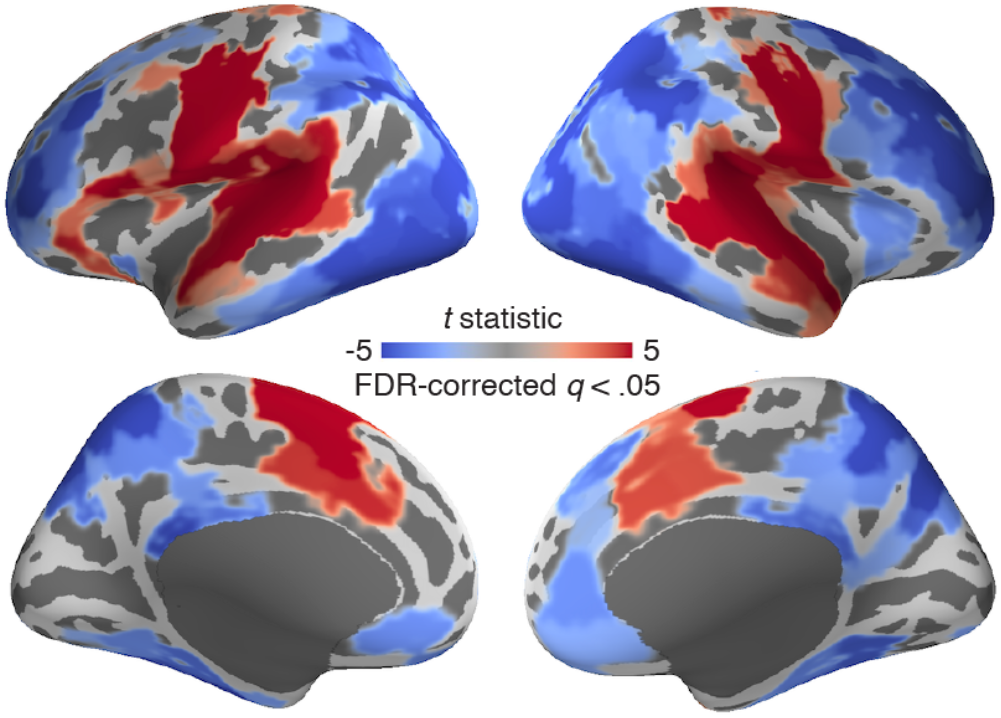
Univariate activation associated with overt speech production. Whole-brain *t*-statistic maps of cortical parcels responding to speech production are displayed on the lateral (top row) and medial (bottom row) surfaces of the inflated fsaverage template brain for both hemispheres. Warmer colors indicate greater activation during speech relative to silence periods, while cooler colors indicate the opposite contrast. Statistical significance was assessed using two-tailed tests (N = 73). Only parcels that survived false discovery rate (FDR) correction across cortical parcels (*q* < 0.05) are shown.

### Comparison between GPT- and human-generated ratings

Overall distributions of participant-level mean GPT-generated ratings across the 14 psychological dimensions are shown in Fig. 4a. Most specific emotions and sensory modalities were reported sporadically, resulting in mean ratings close to 1 (“not at all”).

**Fig. 4.**
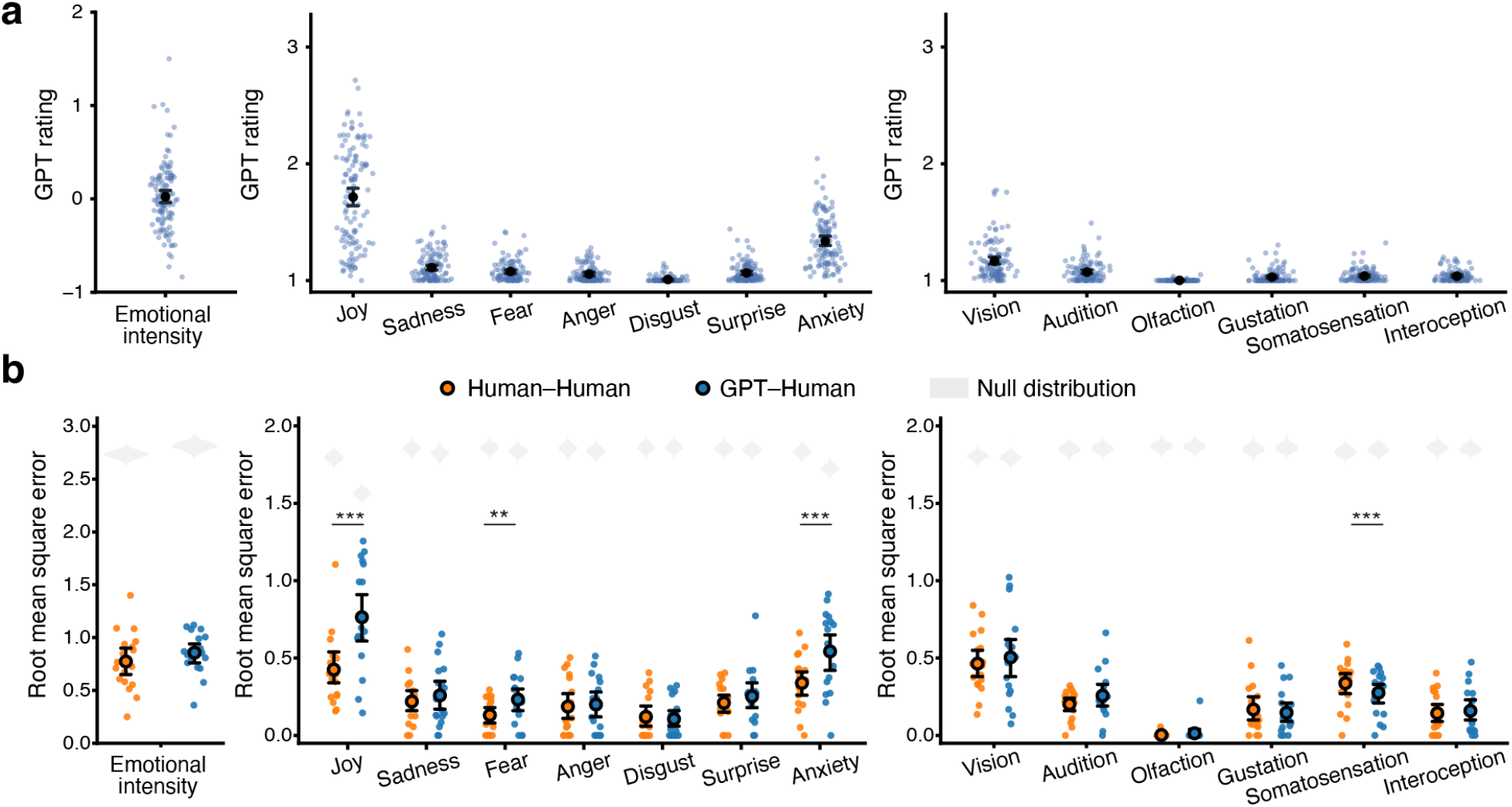
Descriptive statistics and validation of GPT-generated psychological dimension ratings. In both panels, dimensions are grouped into emotional intensity (left), specific emotions (middle), and sensory modalities (right). (**a**) Distributions of the mean GPT-generated ratings. Blue dots represent individual participants’ mean GPT ratings for a given psychological dimension (N = 118), averaged across all thought segments. Black circles indicate the mean across participants, with error bars showing 95% bootstrap confidence intervals. Emotional intensity was rated on a −4 (very negative) to +4 (very positive) scale, whereas all other dimensions were rated on a 1 (not at all) to 4 (very much) scale. (**b**) Validation of GPT-generated ratings through comparison with human ratings. Averaged pairwise root mean square error (RMSE) values were used to quantify rating agreement between human raters (orange) and between GPT and human raters (blue). Colored dots represent selected individual participants (N = 18), larger black-outlined circles indicate the mean across participants, and error bars denote 95% bootstrap confidence intervals. Gray violins show null RMSE distributions generated using random ratings within the valid response range for each dimension. \*\**q* < 0.01, \*\*\**q* < 0.001 (FDR-corrected, two-tailed paired *t*-tests). See Supplementary Table 1 for the full statistics.

To assess the validity of the GPT-generated ratings, we quantified agreement between GPT and human raters for the randomly selected 18 transcripts using root mean square error (RMSE), an absolute error metric that captures the magnitude of disagreement on the original rating scales (Fig. 4b). RMSE can be computed even when there is no variance across ratings (i.e., situations in which correlations are undefined), which makes it well suited for our data where many ratings took the identical value of 1. For each dimension within each transcript, RMSE values were computed between the GPT-generated ratings and each human rater’s ratings, yielding four GPT-Human comparison pairs. These values were then averaged to obtain participant-level GPT-Human RMSE estimates for that dimension. For comparison, participant-level Human-Human RMSE values were also calculated for each dimension by computing RMSE across all possible pairs of the four human raters (six Human-Human pairs) and averaging these values across pairs.

Statistical significance of the group-level mean RMSE was assessed using randomization tests conducted separately for GPT-Human and Human-Human RMSE. For each dimension, a null distribution of mean RMSE representing chance-level agreement was generated using 1000 iterations. In each iteration, ratings were randomly sampled for individual think-aloud sentences within the valid response range of the corresponding dimension for each transcript. For the GPT-Human comparison, RMSE was computed between the randomly generated ratings and the GPT-generated ratings. For the Human-Human comparison, RMSE was computed between the random ratings and each human rater’s ratings and then averaged across raters. The resulting RMSE values were subsequently averaged across all 18 transcripts to obtain a group-level null RMSE for each iteration. Across all 14 dimensions, both GPT-Human and Human-Human ratings showed significantly lower RMSE values than the null distribution (all *p*s < 0.001), indicating that both rating approaches performed substantially better than chance.

We next compared GPT-Human RMSE with Human-Human RMSE to determine if the LLM’s level of agreement with human raters was comparable to the agreement observed among human raters themselves (Fig. 4b). Paired *t*-tests were conducted across participants separately for each dimension, and FDR correction was applied across dimensions to account for multiple comparisons. For the majority of dimensions, GPT-Human RMSE showed levels of agreement comparable to Human-Human RMSE, and even stronger agreement for somatosensation (*t*(17) = 7.12, *q* < 0.001, Cohen’s *d*z = 1.68, 95% CI = [0.04, 0.08]), supporting the validity of the GPT-generated ratings. Exceptions were observed for joy, fear, and anxiety, for which GPT-Human RMSE values were higher than Human-Human RMSE values (FDR-corrected *q*s < 0.05; see Supplementary Table 1 for full statistical results across all dimensions).

### fMRI validation of GPT-generated ratings

We further evaluated the validity of the GPT-generated sensory ratings by testing whether thoughts rated as containing content related to a specific sensory modality (ratings ≥ 2) were associated with increased activation in corresponding perceptual brain regions, relative to thoughts rated as lacking such sensory content (rating = 1). For this analysis, we focused on the two most frequently reported sensory modalities: vision and audition.

After excluding 43 participants based on motion and data quality criteria, participants who produced at least one thought segment (spanning at least 1 TR) rated as containing visual content (N = 72) or auditory content (N = 57) were included in the visual and auditory activation analyses, respectively. For each participant and cortical parcel, preprocessed BOLD signals were averaged across TRs corresponding to thoughts containing the relevant sensory content, and separately across TRs corresponding to thoughts lacking that content. To account for the hemodynamic response delay, timestamps marking the onset and offset of each thought segment were shifted forward by 4.5 s. Motion outlier TRs (FD ≥ 1 mm), along with the two immediately preceding and following TRs, were excluded from averaging. Within-subject contrast values were then computed as the difference between mean activation in the sensory-present and sensory-absent conditions. Group-level effects were assessed using two-tailed one-sample *t*-tests on these contrast values, with multiple comparisons across parcels controlled using the FDR procedure (FDR *q* < 0.05).

The resulting whole-brain group-level contrast maps are shown in Fig. 5. As expected, thought segments rated as containing visual content elicited activation in regions associated with high-level visual perception, including the dorsal parietal and ventral temporal cortices, which are components of the dorsal and ventral visual processing streams^91^ (Fig. 5a). In contrast, thought segments rated as containing auditory content elicited activation in regions associated with auditory perception, including the primary and secondary auditory cortices within the superior temporal lobes (Fig. 5b). These activation patterns provide convergent support for the validity of the GPT-generated ratings in capturing neural responses relevant to thought content.

**Fig. 5.**
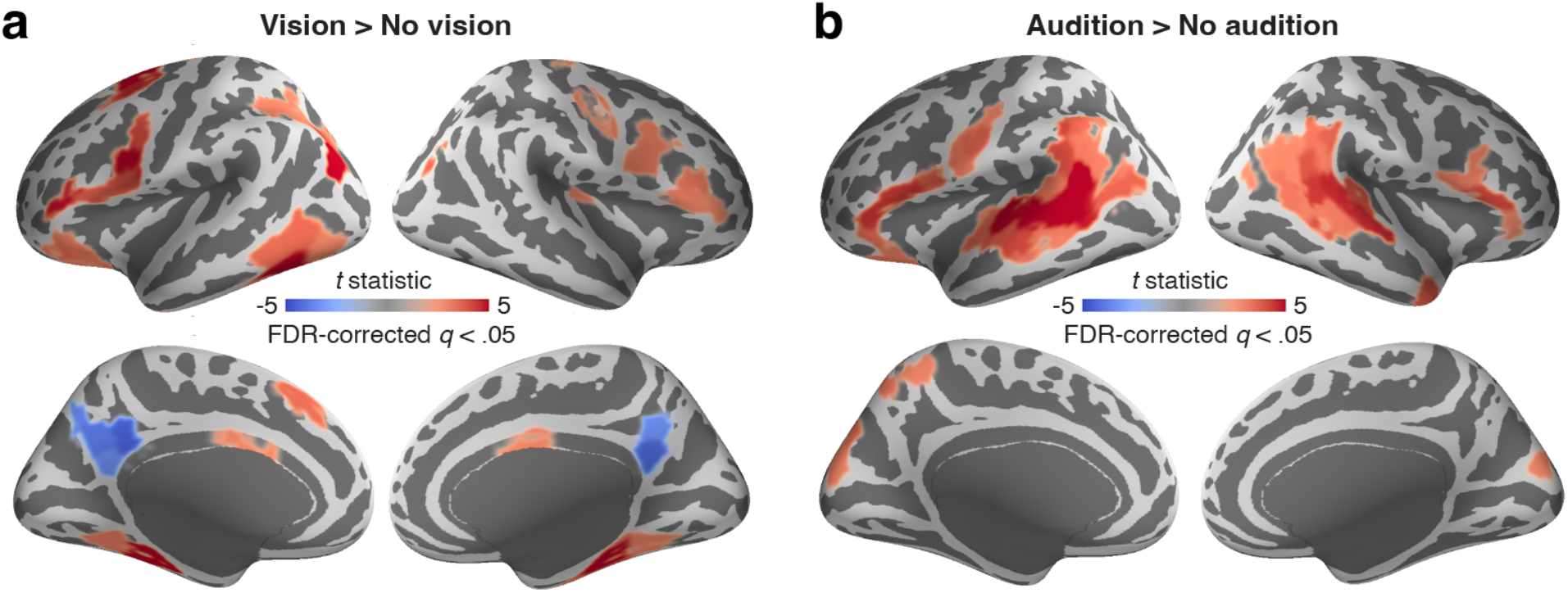
Univariate activation associated with sensory content during think-aloud. (**a**) Whole-brain *t*-statistic maps of cortical parcels showing greater activation during thoughts rated by GPT as containing visual content compared to thoughts rated as lacking visual content (N = 72). (**b**) Whole-brain *t*-statistic maps of cortical parcels showing greater activation during thoughts rated as containing auditory content compared to thoughts rated as lacking auditory content (N = 57). In both panels, *t*-statistic maps are displayed on the lateral (top row) and medial (bottom row) surfaces of the inflated fsaverage template brain for both hemispheres. Parcels showing significantly greater activation, after false discovery rate (FDR) correction across cortical parcels (*q* < 0.05), are shown in red. Statistical significance was assessed using two-tailed tests.

### Post-scan survey validation

To evaluate the quality of the post-scan survey data, we first examined internal consistency across items within each questionnaire. As shown in Table 1, most measures demonstrated acceptable to excellent internal consistency (Cronbach’s *α* > .70)^92,93^, supporting their reliability for individual-differences analyses.

**Table 1.** Descriptive statistics and internal consistency for each survey scale.

| Scale | N items | Mean (SD) | Cronbach's $\alpha$ |
| --- | --- | --- | --- |
| Big5-Openness | 10 | 36.68 (6.82) | 0.809 |
| Big5-Conscientiousness | 9 | 32.87 (5.79) | 0.807 |
| Big5-Extraversion | 8 | 24.06 (6.53) | 0.874 |
| Big5-Agreeableness | 9 | 34.68 (5.27) | 0.773 |
| Big5-Neuroticism | 8 | 23.86 (5.96) | 0.814 |
| SAM-Episodic | 6 | 24.16 (6.81) | 0.831 |
| SAM-Semantic | 6 | 19.01 (4.08) | 0.572 |
| SAM-Spatial | 6 | 20.91 (4.33) | 0.692 |
| SAM-Future | 6 | 22.33 (5.35) | 0.871 |
| Mind wandering (MWQ) | 5 | 18.70 (4.56) | 0.777 |
| Mindfulness (MAAS) | 15 | 3.68 (0.74) | 0.836 |
| Suppression (WBSI) | 15 | 47.26 (11.93) | 0.900 |
| Automatic thought (ATQ-30) | 30 | 51.87 (22.14) | 0.967 |
| ADHD (ASRS v1.1) | 6 | 8.45 (3.58) | 0.622 |
| Narcissism (NPI-40) | 40 | 12.16 (6.66) | 0.853 |
| Self-esteem (RSES) | 10 | 30.25 (4.45) | 0.823 |
| Self-compassion (SCS-SF) | 12 | 3.02 (0.61) | 0.794 |
| Selfishness | 6 | 18.64 (4.37) | 0.746 |
| Curiosity and exploration (CEI-II) | 10 | 34.99 (7.18) | 0.875 |
| DASS-Depression | 7 | 8.12 (7.59) | 0.852 |
| DASS-Anxiety | 7 | 6.23 (6.60) | 0.787 |
| DASS-Stress | 7 | 11.16 (7.66) | 0.775 |
| PTSD (PCL-5) | 20 | 12.61 (12.11) | 0.922 |
| Trait anxiety (STICSA) | 21 | 31.41 (8.39) | 0.879 |
| Social phobia (SPIN) | 17 | 17.93 (13.97) | 0.936 |
| Rumination (RRS-10) | 10 | 20.49 (6.30) | 0.844 |
| Narrative engageability | 16 | 4.38 (1.30) | 0.944 |

To further assess construct validity, we examined patterns of association among conceptually related constructs by computing correlations across survey measures (Supplementary Fig. 1). As expected, measures indexing related psychological constructs were correlated in theoretically consistent directions. For example, depressive symptoms measured by the DASS-21 were strongly correlated with rumination from RRS-10 (*r*(67) = 0.59, *p* < 0.001) and automatic negative thoughts from the ATQ-30 (*r*(67) = 0.71, *p* < 0.001). PTSD symptoms were also positively associated with depression (*r*(67) = 0.57, *p* < 0.001), anxiety (*r*(67) = 0.53, *p* < 0.001) and automatic negative thoughts (*r*(67)= 0.63, *p* < 0.001). The three DASS-21 subscales (depression, anxiety, and stress) were strongly intercorrelated (*r*s ≈ 0.58–0.68, *p* < 0.001). In addition, mind wandering from the MWQ was strongly negatively associated with mindfulness from the MAAS (*r*(67) = -0.57, *p* < 0.001), in line with previous research^94^. These patterns provide support for the construct validity of the survey responses.

## DATA AVAILABILITY

Neuroimaging data are available through the OpenNeuro repository (accession number: ds006067; version 2.0.0)^72^. Behavioral and annotation data are available via the Open Science Framework (https://osf.io/a56rm)^73^, including think-aloud transcripts with word-level timestamps, sentence-level psychological dimension ratings, and post-scan survey responses.

## CODE AVAILABILITY

Scripts used for MRI data preprocessing, timestamp and rating generation, and validation analyses are publicly available via the Open Science Framework, in the “code” directory of the “Think Aloud Behavioral Data” project (https://osf.io/a56rm)^73^.

## AUTHOR CONTRIBUTIONS

H.L. conceived and designed the research. M.Zhang, P.R.L, X.L., S.B., Y.L, J.C. and H.L. collected the data. M.Zhang, P.R.L, H.S, and H.L. analyzed the data. M.Zhang, P.R.L, and H.L. wrote the original manuscript. M.Zhang, P.R.L, H.S., M.Zhao, X.L., C.J.H., and H.L. reviewed and edited the manuscript. C.J.H., J.C. and H.L. provided funding.

## ACKNOWLEDGEMENTS

We thank Wenyu Chen for assistance in generating word-level timestamps.

## SUPPLEMENTARY FIGURE

**Supplementary Figure 1.**
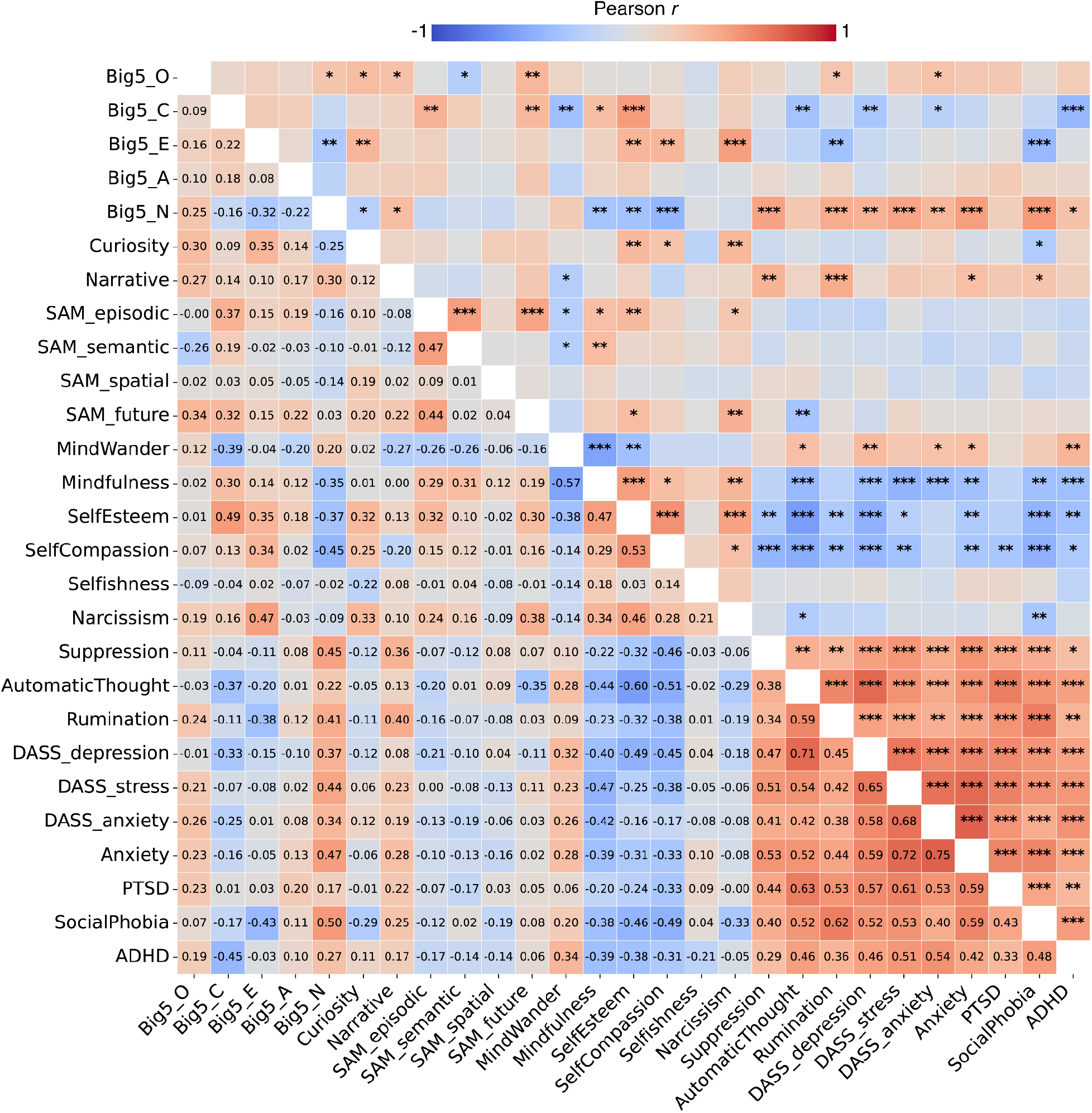
Correlations among survey scores (*N* = 69). The lower triangle of the heatmap displays pairwise Pearson correlation coefficients (*r*) for all questionnaire measures. The upper triangle shows statistical significance markers (\**p* < 0.05, \*\**p* < 0.01, \*\*\**p* < 0.001, uncorrected). Warm colors indicate positive correlations and cool colors indicate negative correlations.

## SUPPLEMENTARY TABLE

**Supplementary Table 1.** Comparison of the root mean square error (RMSE) across human raters and between human raters and GPT-5 for each psychological dimension.

| Dimension | H-H mean (SD) | G-H mean (SD) | *t*(17) | 95% CI | Cohen's *d**z* | *p* | FDR *q* |
| --- | --- | --- | --- | --- | --- | --- | --- |
| Emotional intensity | 0.77 (0.27) | 0.86 (0.20) | 1.68 | [-0.02, 0.19] | 0.40 | 0.111 | 0.259 |
| Joy | 0.43 (0.23) | 0.76 (0.33) | 4.82 | [0.19, 0.49] | 1.14 | < .001 | < .001 |
| Sadness | 0.22 (0.14) | 0.26 (0.20) | 1.42 | [-0.02, 0.10] | 0.33 | 0.175 | 0.272 |
| Fear | 0.13 (0.11) | 0.23 (0.16) | 3.97 | [0.05, 0.15] | 0.94 | < .001 | 0.003 |
| Anger | 0.19 (0.18) | 0.20 (0.17) | 0.88 | [-0.02, 0.05] | 0.21 | 0.394 | 0.431 |
| Disgust | 0.12 (0.14) | 0.11 (0.12) | -1.19 | [-0.04, 0.01] | -0.28 | 0.251 | 0.319 |
| Surprise | 0.21 (0.13) | 0.25 (0.18) | 1.43 | [-0.02, 0.10] | 0.34 | 0.172 | 0.272 |
| Anxiety | 0.34 (0.18) | 0.54 (0.26) | 4.70 | [0.11, 0.30] | 1.11 | < .001 | < .001 |
| Vision | 0.46 (0.19) | 0.50 (0.28) | 1.22 | [-0.03, 0.11] | 0.29 | 0.241 | 0.319 |
| Audition | 0.20 (0.09) | 0.26 (0.16) | 2.21 | [0.00, 0.11] | 0.52 | 0.041 | 0.116 |
| Olfaction | 0.00 (0.01) | 0.01 (0.05) | 0.86 | [-0.02, 0.04] | 0.20 | 0.401 | 0.431 |
| Gustation | 0.17 (0.17) | 0.15 (0.14) | -1.45 | [-0.05, 0.01] | -0.34 | 0.164 | 0.272 |
| Somatosensation | 0.34 (0.15) | 0.28 (0.14) | -7.12 | [-0.08, -0.04] | -1.68 | < .001 | < .001 |
| Interoception | 0.14 (0.13) | 0.16 (0.15) | 0.70 | [-0.03, 0.06] | 0.16 | 0.495 | 0.495 |

*Note:* H-H = Human-Human RMSE (across human raters). G-H = GPT-5-Human RMSE (across GPT-5 and human raters). 95% CI = 95% confidence interval for the difference between G-H RMSE and H-H RMSE. Cohen's *d**z* = standardized mean difference for paired samples. *p*-values are uncorrected, and FDR-corrected *q* values are reported to account for multiple comparisons across dimensions.

